# A genomic view of the peopling of the americas

**DOI:** 10.1101/058966

**Authors:** Pontus Skoglund, David Reich

## Abstract

Whole-genome studies have documented that most Native American ancestry stems from a single population that diversified within the continent more than twelve thousand years ago. However, this shared ancestry hides a more complex history whereby at least four distinct streams of Eurasian migration have contributed to present-day and prehistoric Native American populations. Whole genome studies enhanced by technological breakthroughs in ancient DNA now provide evidence of a sequence of events involving initial migration from a structured Northeast Asian source population, followed by a divergence into northern and southern Native American lineages. During the Holocene, new migrations from Asia introduced the Saqqaq/Dorset Paleoeskimo population to the North American Arctic ~4,500 years ago, ancestry that is potentially connected with ancestry found in Athabaskan-speakers today. This was then followed by a major new population turnover in the high Arctic involving Thule-related peoples who are the ancestors of present-day Inuit. We highlight several open questions that could be addressed through future genomic research.

## Introduction

The peopling of the Americas represented the culmination of a Late Pleistocene expansion of anatomically modern humans out of Africa. Archaeological evidence indicates that groups subsisting on hunting lived in extreme northeast Siberia (71°N) by at least 28,000 years ago [1]. Human groups adapted to the mammoth steppe habitat were thus poised to enter Beringia—the landmass between Alaska and Eurasia that is now submerged—by this time [2,3]. The path from Beringia to the more temperate parts of the American continents, however, was blocked by the merged Laurentide and Cordilleran ice sheets that covered northern North America until the end of the Last Glacial Maximum. The ice retreated from parts of the Pacific coast ~16,000 years ago, raising the possibility of a coastal migration after this time, and within a few thousand years a habitable corridor through the center of the continent opened between the two ice sheets [4]. The first unambiguous evidence of modern humans in the Americas dates to between 14,000 and 15,0 years ago [5–8], and was likely the consequence of migration from Beringia.

Major debates about the peopling of the Americas have focused on the question of whether the first early human populations in the Americas are directly ancestral to present-day Native Americans, as well as on the number, mode, and timing of secondary migrations to the Americas. Advances in genomics have, within the last five years, enabled the collection of far more data from present-day Native Americans than was available previously, and have also made it possible for the first time to access DNA from ancient Native American remains. Analysis of these data has highlighted a minimum of four distinct streams of Asian ancestry some of which were not clear from the archaeological evidence. We review the contributions of genomic data to understanding the prehistory of the Americas, and highlight outline outstanding questions where it may be able to provide additional insight.

## The power of the whole genome

The first meaningful genetic insights about Native American population history came from mitochondrial DNA, a segment of about 16,500 base pairs (approximately 1/200,000^th^ of the genome) that is passed exclusively along the maternal line. Mitochondrial DNA was one of the first parts of the genome to be heavily investigated to learn about human population history for several reasons. First, it is highly variable on a per-nucleotide level and thus sequencing only a short stretch can detect non-trivial amounts of human variation. Mitochondrial DNA’s high variability and short length meant that it was practical to sequence in large numbers of samples at a time when it was prohibitively expensive to generate genome scale data. For ancient DNA studies, mitochondrial DNA had the further advantage that it exists in about a thousand-fold higher copy number than any other single place in the genome. Since one of the main challenges of ancient DNA is obtaining sufficient amount of material from any position in the genome to be able to analyze, beginning with more starting material can be an advantage [9].

The greatest contribution of mitochondrial DNA studies to the understanding of Native American prehistory has been in the area of reconstruction of population size history. Mitochondrial DNA analyses were the first to document that the ancestry of most Native Americans derives from a population that experienced a profound founder event [10], with a relatively small number of individuals giving rise to a large number of descendants today. The evidence for this is that all Native American mitochondrial DNA lineages today descend from just five founding maternal lines [11–13] that each had a common ancestor around 18,000 to 15,0 years ago, implying a population size bottleneck around this time [14–18]. The evidence for a profound population bottleneck has since been confirmed and its intensity measured more accurately with genome scale data [19–23], but it is important to note that there are still challenges with disentangling the number of founder individuals from the duration of the population size reduction using all the reported methods.

A second finding about Native American population history based on mitochondrial DNA data is the evidence that the founder event may have been proceeded by an extended period (many thousands of years) of little or no shared ancestry with non-Native American mitochondrial DNA lineages. This suggested to some researchers the hypothesis of a ‘Beringian standstill’, whereby the first founding population of the Americas was isolated from Eurasian populations prior to its radiation into a multitude of sub-populations in America [15].

At the same time, some observations from mitochondrial DNA studies of the Americas have been more confusing than helpful. For example, the mitochondrial DNA subtype called D4h3a is today almost entirely restricted to Pacific coastal populations, both in North and South America. This observation led to the hypothesis that D4h3a was carried by the members of a population that carried Native Americans south of the ice sheets along a coastal route, in a migratory movement that was distinctive from what led to many other Native American populations [24]. However, ancient DNA studies have since found the same mitochondrial DNA type in a ~12,600 year old individual from present-day Montana, which based on its genome-wide data is unambiguously from the main ancestral lineage leading to most Native Americans [25].

It is now clear that so many founder events and fluctuations in population size have occurred before, during, and after the peopling of the Americas that the evidence from one position in the genome—mitochondrial DNA, the Y chromosome, or any other location—is too subject to random changes in frequency (genetic drift) to be meaningful by itself. Only by taking the independent testimony of many locations in the genome simultaneously can we obtain a high-resolution picture of the deep past. The remainder of this article focuses on insights from whole genome studies about Native American history. While these studies are still in their early days, they have already upended our understanding of key events in Native American population history. Application of ancient DNA technology promises further insights in years to come.

## Sources of Native American ancestry

Under the hypothesis where Native American ancestry stems from a single founder population that separated earlier from Eurasian populations, differences in allele frequencies between Native American groups should have developed independently from Eurasian allele frequencies. This simply null hypothesis makes it possible to explicitly test hypotheses about the number of American founder populations. Reich *et al.* [23] applied this idea to the first comprehensive genome-wide data from Native American populations (52 populations, but none from the continental United States), and concluded that at least three ancestral populations—or streams of gene flow—were required to explain the similarities between Native Americans and East Asians. According to the initial study [23], all Native American groups from Central and South America fit a model of a single founder population. An additional source of ancestry was necessary to explain genetic variation in Eskimo-Aleut speakers. In addition, analysis of the Athabaskan-speaking Chipewyan revealed that they could not solely have their ancestry from the same founding population as other Northern-, Meso-and South American populations.

## The main ancestral stream giving rise to Native American ancestry

One of the most important pieces of genetic evidence relevant to the peopling of the Americas was the sequencing of a genome from the remains of a child (‘Anzick-1’) buried with Clovis artifacts in western Montana and directly dated to 12,600 before present (BP) [25]. This child was consistent with deriving all of his ancestry from the same founding population as Central and South Americans (Figure 1), contradicting the ‘Solutrean hypothesis’ [26] that posits genetic discontinuity between the makers of the Clovis industry and present-day Native Americans [27].

**Figure 1.**
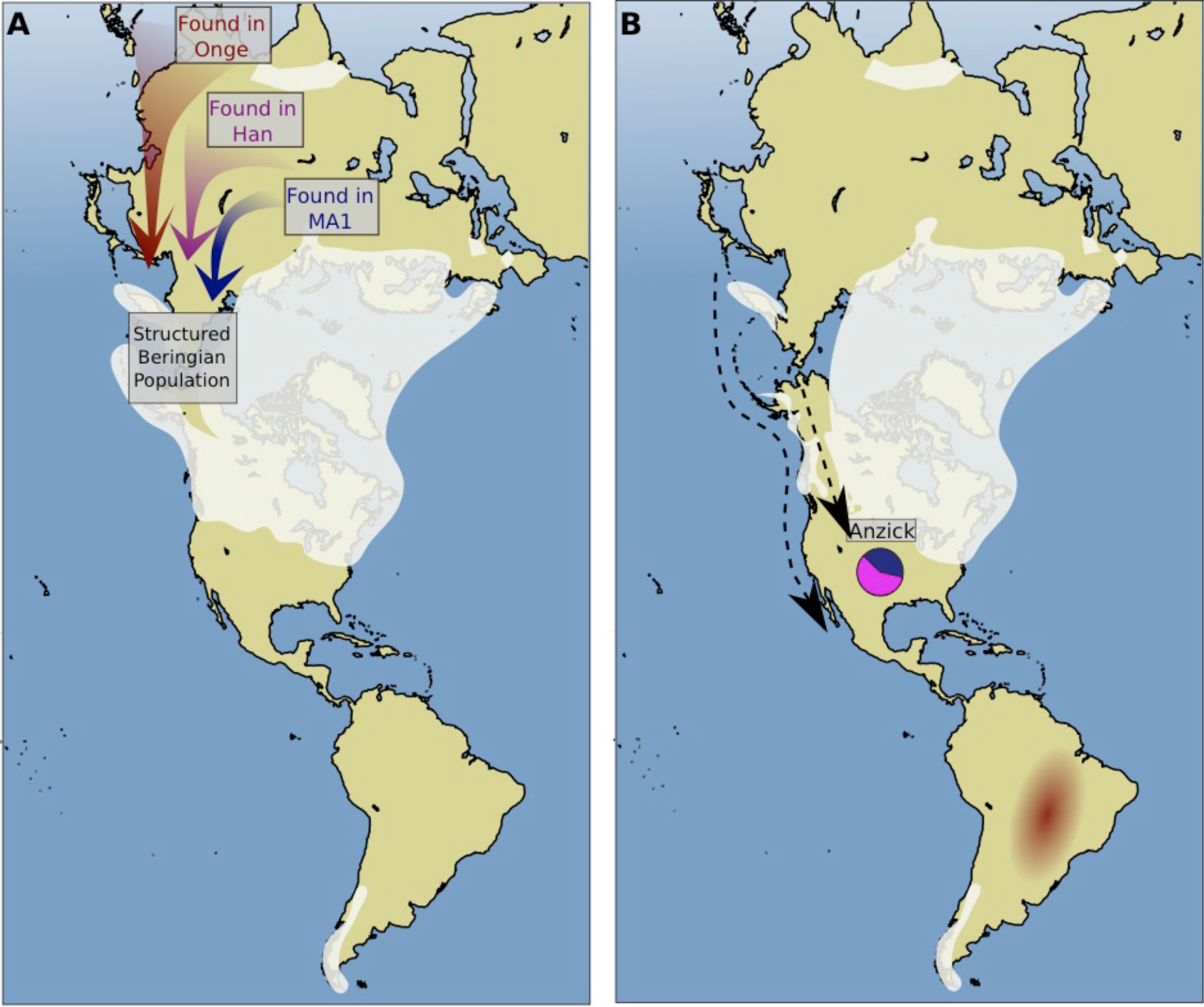
Eurasian source populations of the earliest Native Americans. A) Diverse population lineages that genetic evidence suggest were present in Northeast Asia and contributed to the ancestry of present-day Native Americans. These comprise people with ancestry related to present-day Andamanese and Australo-Melanesians (red), people related to present-day East Asians such as Han Chinese (pink), and people related to the 24,000 year old Mal’ta child buried near Lake Baikal (dark blue). By 12,600 years ago, at least the Mal’ta-related and east Asian-related ancestries were already present in the proportions found in Native Americans today in the Anzick child from Western Montana (B). Today, the Andamanese-related ancestry can be detected as an excess affinity to present-day Amazonians (B).

The most surprising finding was that the Anzick individual is from a population more closely related to Central-and South Americans than to some northern North Americans (including all speakers of Algonquian languages studied to date), despite the apparent common ancestral origin of Native Americans across the continents. This suggests that the present-day population structure of the main ancestry in Native Americans [23] dates back to more than 12,600 years ago [25], and that this diversification divided the ancestry of present-day Native Americans into two main streams, one of which includes the ancestors of present-day Northern Native Americans analyzed (‘NNA’: Cree, Ojibwa, and Algonquin), and the other of which includes the Anzick individual and present-day Central-and South American groups (‘SA’:*e.g.* Mixe, Quechua, and Yaghan).

While we thus have evidence for SA ancestry in the Late Pleistocene in the form of the Anzick genome, an outstanding question is where the ancestors of NNA were localized. One possibility is that they were confined to ice-free regions north of the corridor between the Cordilleran and Laurentide ice sheets, and only expanded south after 12,600 BP, where this ancestry would eventually displace the SA populations represented by Anzick (Figure 2). Another possibility might be an expansion of the Clovis industry from a southern origin that also represented a population expansion by SA populations into regions where NNA populations were located (Figure 2). Thus SA populations might have brought Clovis technology to some regions in northern North America only to later be displaced by an NNA resurgence.

**Figure 2.**
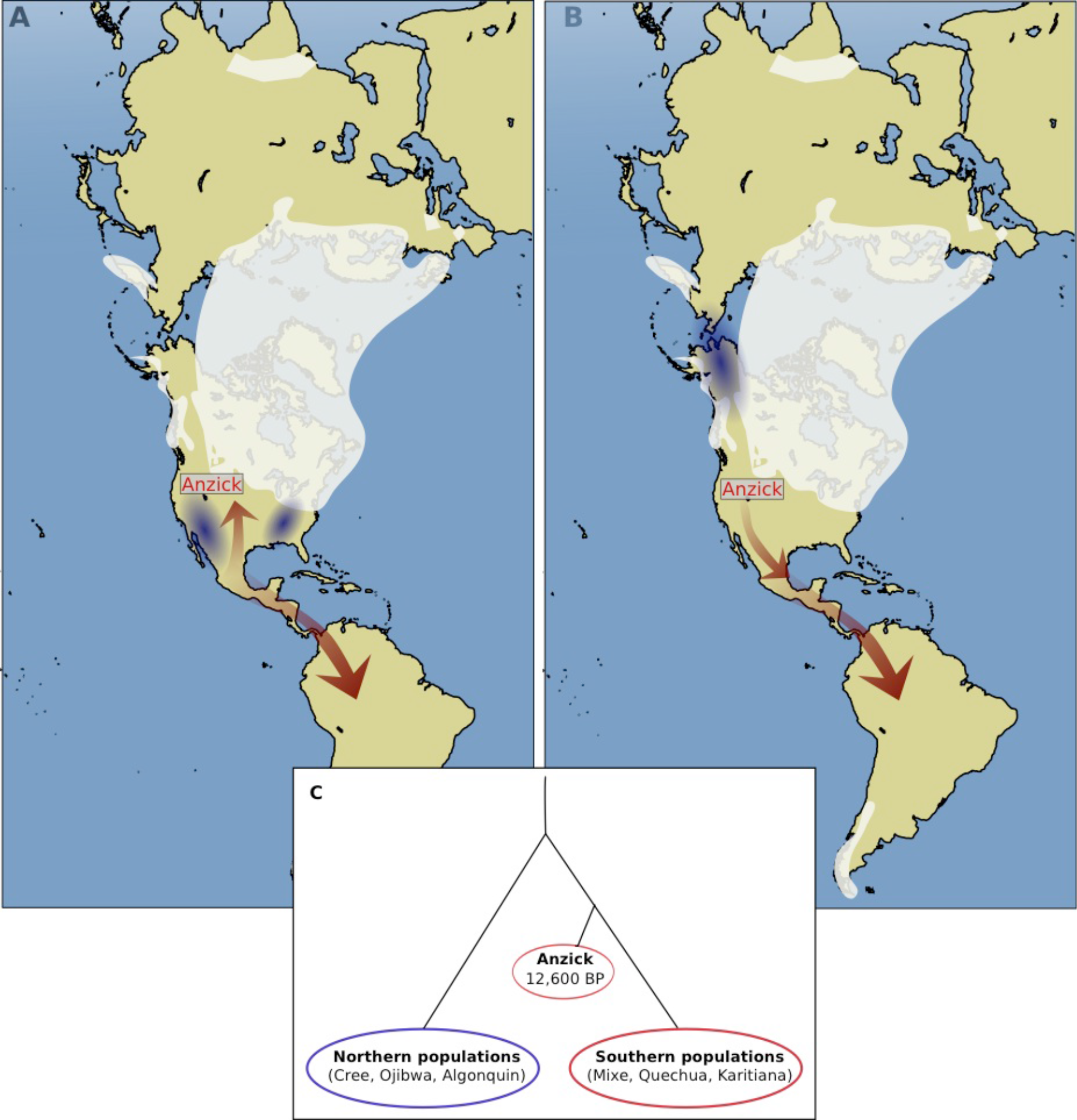
Two scenarios explaining that the 12,600 year old Anzick individual is more closely related to South and Central-Americans than Northern Native Americans. A) Clovis technology appears first in the southern parts of North America. It is thus possible that Anzick, being associated with Clovis artifacts and dating to the end of the Clovis period, represents a northwards expansion into a region where the ancestors of present-day northern Native Americans were already established. B) An alternative is that the southern lineage associated with the Anzick individual represents the first migration south of the ice sheets, whereas the ancestors of present-day Northern Native Americans resided north of the ice sheets at 12,600 years ago and only later migrated southwards, replacing the Anzick-like population. C) Both these scenarios could explain the observation that Anzick is more closely related to Southern Native American populations.

More recent genome sequencing [28] of the ~9,000 year-old Kennewick individual (‘the Ancient One’) did not provide resolution about this issue. While the genome analysis showed that the Kennewick individual had the great majority of its ancestry from the same deep source as other present-day Native Americans, his affinity to the SA and NNA lineages remains ambiguous.

## Major admixture during the formation of the ancestral population of Native Americans

Until 2014, efforts to infer demographic parameters for the peopling of the New World based on genetic data focused on modeling Native Americans as an offshoot of East Asian ancestry [17,20,23,29–34]. However, the analysis of the genome of a 24,000 year-old individual from the Mal’ta site near Lake Baikal in Central Siberia revealed that this model is untenable. The Mal’ta individual shared genetic affinities to both European (West Eurasian) and Native American populations [35]. Analyses showed that a scenario in which Mal’ta descended from an admixture between a West Eurasian population and an ancient population that was also ancestral to Native Americans could not explain all features of the data [35]. However, a scenario in which Native Americans are admixed between lineages related to Mal’ta (between 1/4 and 1/2 of Native American ancestry) and East Asians can explain the data (Figure 1). Thus, Native Americans and East Asians do not in fact descend from a common ancestral population that separated earlier from a lineage leading to Mal’ta and to West Eurasians [35].

The finding of ancient mixture in the ancestry of Native American—prior to diversification within the Americas—also has consequences for modeling other features of Native American population history. Genetic debate about the date of the first migrants into the Americas has had, as one of its important themes, estimation of the date of genetic divergence of the lineages giving rise to Native American and East Asian populations [31]. The major admixture related to the Mal’ta lineage in the ancestry of all Native Americans is inconsistent with the assumption of a simple population split between Native Americans and East Asians that has been the basis for the most attempts to date to infer the population split times, and which have suggested dates of around 23,000 years ago [33,34]. The substantial contribution of the Mal’ta lineage to Native Americans may have the effect of upwardly biasing estimates of the time of divergence of Native Americans and East Asians. Future models estimating parameters for the founding populations of the Americas will need to consider this admixture explicitly.

## An Australasian Connection

Recently, we carried out a stringent test of the null hypothesis of a single founding population of Central and South Americans using genome-wide data from diverse Native Americans [36]. We detected a statistically clear signal linking Native Americans in the Amazonian region of Brazil to present-day Australo-Melanesians and Andaman Islanders (‘Australasians’). Specifically, we found that Australasians share significantly more genetic variants with some Amazonian populations—including ones speaking Tupi languages—than they do with other Native Americans. We called this putative ancient Native American lineage “Population Y” after *Ypykuéra,* which means ‘ancestor’ in the Tupi language family.

To learn more about the Population Y ancestry present in the Americas, we carried out a series of statistical modeling analyses. The genetic patterns could be explained by as little as 2% admixture from an Australasian-related population that penetrated deep inside the Americas without mixing with the main ancestral lineage of present-day Native Americans. Alternatively, the patterns could be more explained by a larger proportion of ancestry (2-85%) from a population that existed in a substructured Northeast Asia, and was similar to the main lineage that gave rise to other Native Americans while retaining more Australasian affinity.

We considered the possibility that these genetic patterns relate to the claims based on skeletal morphology that there was an wave of migration related to Australasians that contributed to early Native Americans, followed by later large-scale population replacement by the primary ancestors of present-day Native Americans [37,38]. While this evidence has been contested on morphological grounds—with the most important critique being that the analyses are not statistically compelling [2,39]—the fact that the morphological evidence is strongest in Brazil where Population Y ancestry is prevalent justifies investigation into possible connections. However, new genetic findings reject one set of the arguments based on morphology: that groups such as historic-period Native Americans in the Baja California region of Mexico and the Tierra del Fuego region in the southern tip of South America are related to the hypothesized earlier Native American migration. Genomic DNA from these populations do not have any evidence of affinity to Australasian populations [34].

What is the history behind the Population Y ancestry in Amazonian Native Americans today? Little can be said at present, as we do not have ancient DNA from individuals carrying detectable amounts of this ancestry. The implication that the Siberian populations that gave rise to the ancestors of Native Americans was substructured, however, is not surprising given that the 24,000-18,000 years from the Lake Baikal region of Siberia (the Mal’ta and Afontova Gora sites) were genetically very divergent from present-day East Asians [35,40]. The Population Y results suggest that a population that has not yet been sampled with ancient DNA data—one with more Australasian-related ancestry—may also have been present in the broad geographic area to contribute to the founders of Native American founders. Notably, Andaman Islanders, the population with the single strongest affinity to Amazonians, are not as good match for the non-Marta like ancestry in Central Americans as are Chinese populations [36]. These strands of evidence suggest a minimum three-part ancestry of the Beringian populations that came to populate the Americas (Figure 1). Two of these strands were fully braided together to form the main ancestral lineage of Native Americans by time of the Beringian bottleneck. However, the third strand, with an affinity to Australasians, was not.

It has been suggested that Native American ancestors may have entered the more temperate parts of North American both by an early coastal route and the later ice free corridor [2]. One possibility is that the different sources of deep ancestry that are inferred by these genetic patterns may reflect movements of a substructured Beringian population through these different routes. Alternatively, the patterns could reflect pulses of migration from Beringia occurring at different times (e.g. multiple pulses through the ice free corridor). Restricting study to South America, a related question is the history behind the deeply structured population lineages East and West of the Andes in South America that have been documented based on genomic data [23]. Population Y ancestry may be limited entirely to the eastern populations, raising the possibility that this split was extremely ancient.

**Figure 3.**
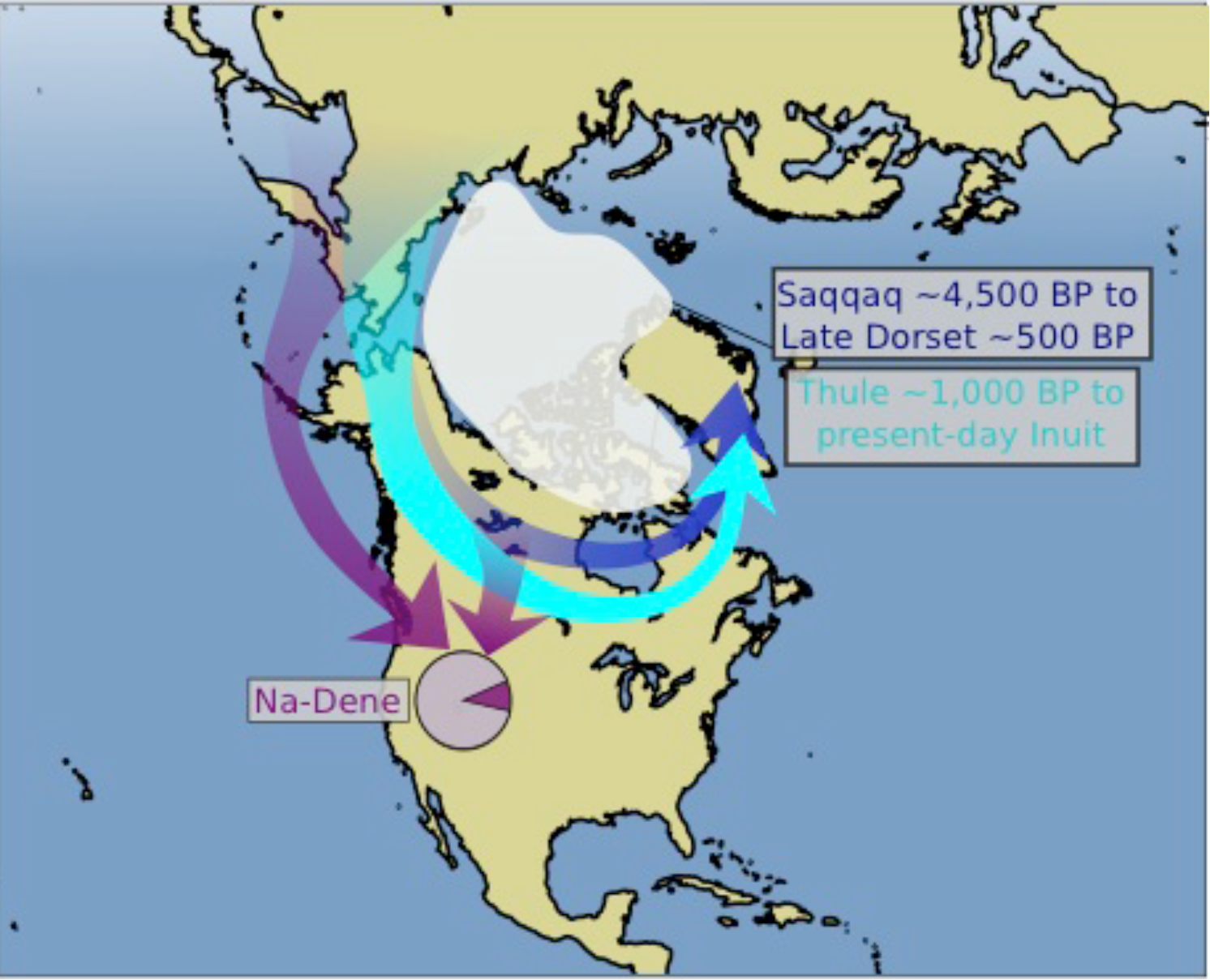
Holocene migrations to the Americas. There is a current consensus supporting a Paleoeskimo migration to Greenland ~4,500 years ago, giving rise to both the Saqqaq culture and the Early, Middle and Late Dorset cultures. This Paleoeskimo population was displaced by Neo-Eskimos associated with the Thule culture migrating through the Arctic ~1,000 years ago, although there remains a possibility of some admixture. Two scenarios can plausibly explain the Asian ancestry flow in present-day speakers of Athabaskan (Na-Dene) languages. 1) The Na-Dene have ancestry from a distinct migration to the Americas from that seen in any other ancient samples. 2) The Na-Dene-specific ancestry comes from the same migration as the Paleoeskimos [23]. A proposed model in which the Na-Dene-specific ancestry is from the same migration that contributed to the Inuit [34] has been rejected statistically [23].

## Post-glacial migrations into North America

The archaeological record of the North American Arctic documents a transformation beginning around a thousand years ago and associated with the Thule cultural complex. The Thule culture advanced into a North American Arctic occupied by the descendants of previous migrations, referred to as Paleoeskimos and culturally represented by the Saqqaq and Dorset technological complexes [41]. Genome-wide data from a ~4,000 year old Saqqaq culture individual in Greenland—the first genome-wide dataset from an ancient human—was interpreted as showing that the Saqqaq population went extinct in North America, since the Saqqaq genome was most similar to present-day Koryak and Chukchi on the eastern side of the Bering strait in clustering analyses [42]. Genome-wide analysis of additional samples from the Early Dorset and Middle Dorset cultures—along with Thule individuals—confirmed the picture of distinct ancestry in Paleoeskimos, and a population break associated with the Thule that led directly to present-day Inuit populations [43].

A major outstanding issue in the interpretation of the peopling of the Arctic relates to the geographic and temporal origins of the distinct Asian ancestry present in speakers of Athabaskan (Na-Dene) languages [23,43,44]. Two distinct models for Paleoeskimo and Athabaskan-speaking ancestry have been proposed based on the genetic data. Raghavan *et al.* [43] proposed that Athabaskan speakers do not harbor ancestry from Paleoeskimos; instead, they suggest that the east Asian affinities in the Athabaskans may be due to admixture with Inuit (Neo-Eskimos). Reich *et al.* [23] reported a statistical test that significantly rejects this hypothesis: they showed that ancestry in Athabaskan speakers cannot be modeled as a mixture of the lineages that have given rise to other Native American and Inuit population living today. However, a model in which Athabaskan-speakers harbor about 10% of their ancestry from a source that is distinct from both the Inuit and the main ancestry in other Native Americans—possibly related to the Saqqaq Paleoeskimo—may be consistent with the data [23]. However, several of these inferences may be complicated if Siberian populations today have substantial amounts of ancestry from back-migrations out of America. An important direction for future research is to determine the origin of the distinct Asian ancestry present in Athabaskans using higher resolution methods and additional data.

## Prospects

Far more remains to learn about the genomic history of Native Americans than has already been discovered. Major directions for future data collection include more ancient DNA data, and filling in sampling gaps of present-day Native American populations, especially in United States. We conclude by highlighting five major open questions about Native American population history that we believe may be meaningfully addressed using genomic data in the coming years.

(1) What is the origin of Population Y in Amazonia - can we find it in ancient DNA?
(2) What is the origin of the lineage found in many present-day northern North Americans but not in the 12,600 year old Anzick genome?
(3) What is the history of peopling and migration in South America? Was there an early population split east and west of the Andres, and how much major migration occurred after the first people arrived?
(4) What is the genetic legacy of the Paleoeskimos? Can this widespread population really have disappeared completely after the arrival of the Thule and Aleuts or have they left some descendants, perhaps in admixed form in Athabaskan speakers?
(5) What was the structure of Native American populations in North America prior to the disruption of the last 500 years?

We conclude by emphasizing that true understanding of the population history of any group or region cannot be achieved through genomic studies alone, but requires a synthesis of insights from genomics with information from anthropology, linguistics, archaeology, and sociology. It will also be important to involve Native American organizations and communities in dialogues about these studies, as their perspectives have been underrepresented in these studies in the past.

## Acknowledgements

We thank Lars Fehren-Schmitz and David Meltzer for critical comments. P.S. was supported by the Wenner-Gren foundation and the Swedish Research Council (VR grant 2014-453). D.R. was supported by NIH grant GM100233, by NSF HOMINID BCS-1032255, and is a Howard Hughes Medical Institute investigator.

